# The stab resistance of *Bombyx mori* silk cocoons

**DOI:** 10.1101/2024.11.10.622846

**Authors:** Ateeq Ur Rehman, Vasileios Koutsos, Parvez Alam

## Abstract

This paper considers the mechanical response of *Bombyx mori* silk cocoons to knife stabbing, a simple but controlled way of simulating predaceous penetration. Here, we stab test both entire cocoons (EC) and cocoon wall segments (CWS) statically and dynamically, and note that the process can be broken down in three stages. The first stage involves material deflection, the second is knife penetration, and the third is knife perforation. We find that ca. 95 % of the kinetic energy is lost during the penetration stage. There are noticeable differences in strain between the equatorial 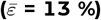 and meridional 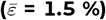 directions before and after the stabbing of EC specimens (p < 0.001). The apparent area of the cocoon is noted to be on average 7 % lower after stabbing than it is prior to being stabbed (p < 0.01). We find that while compression of the cocoon from stabbing results in equatorial expansion (with a Poisson’s ratio, *ν* = 0.25), in the meridional direction the cocoon contracts (*ν* = −0.05) thus showing auxetic behaviour. Force-deflection curves are different in CWS specimens as compared to EC specimens, and this is attributable to natural curvatures in CWS specimens remaining even after a being flattened for mounting and testing. Differences between EC and CWS specimens are also noticeable in the sizes of the stab footprints, with EC samples exhibiting 33 % smaller footprints than CWS samples (p < 0.001). We conclude that testing whole cocoon structures provides a more accurate understanding of their properties as compared to cut and flattened structures. This is because flattening cocoon wall specimens induces delamination and multiple failure zones, reducing the natural stab resistance of the material.

## INTRODUCTION

*Bombyx mori* silk cocoons are characteristically nonwoven composite structures formed into shapes closely resembling ellipsoids, the meridional and equatorial axes being the longer and shorter axes, respectively ^1,2^. The cocoon wall is a hierarchically graded structure comprising numerous layers ^3–6^. The outermost layer of the cocoon has a rough surface with irregular wrinkles. This is understood to be because the silkworm spins the outermost layer first, thus exposing it to the external environment. The outer layer dries quickly in sunlight, resulting in higher levels of shrinkage. In contrast, the inner layers dry more slowly, experiencing lower levels of shrinkage, and exhibiting smoother surfaces as compared to the outermost layer ^7–9^. According to Zhao et al. ^10,11^, the cocoon wall exhibits mechanical anisotropy with graded-layer properties. The cocoon has been studied for its tensile properties ^5,10,12,13^, as well as for moisture management, mass, and heat transfer characteristics ^14–16^. Wang et al. ^17^ report that the outermost layer has the highest puncture strength due to the presence of wrinkles, an elevated sericin fraction, and the highest peel strength when compared against any other layer of the *Bombyx mori* cocoon. They further asserted that wrinkles present on the outer surface enhance frictional resistance, contributing to the puncture resistance of the silk cocoon ^9^.

Cocoons are constructed by moth larvae (pupae), and serve as protection of the pupa from predatorial attack, and environmental hazards ^1,6,18–21^. Silkworms are vulnerable to physical attacks from harsh weather conditions and animals like birds, insects ^22,23^, and rodents ^24^. Parasitic predators such as *Ichneumonid* wasps are known to drill through the cocoon ^23^, while mice and other small mammals use their teeth to puncture through and tear the cocoon wall to reach pupae ^24^. Finally, birds such as *woodpeckers* use their beaks to create holes in the cocoon ^25^. In our previous work, we discovered that the *Bombyx mori* cocoon wall exhibits tear-resistant characteristics, with the outermost layers being more resistant to tearing compared to the innermost ones ^2^. Building on this, we aim to further understand the damage tolerance of this naturally bioengineered structure. Wang et al. ^26^ studied the damage mechanism of the *Bombyx mori* cocoon by needle puncture tests ^9,17^. Similarly, Chen et al. indented the cocoon with a 1.56 mm diameter point indenter to characterise its damage characteristics. Each of these are useful indicators to damage tolerance, however, the teeth of predators such as mice, are known to have sharp, bevelled edges ^27,28^ and as such, teeth like these would not only puncture the cocoon, but would also cut through the cocoon post-puncture. To provide a suitable quantitative assessment of damage tolerance that includes perforation, we propose the use of a knife stab test, as it enables the mimicry of the action of teeth or claws used by predators such as small mammals ^24^. We employ two testing techniques to establish stab resistance from a standard HOSDB knife ^29^. Given that the cocoon protects the larvae during its pupal stage, the cocoon would presumably also have both penetration and perforation resistant properties.

Stab resistance has primarily been researched by textile engineers. The stab-resistant characteristics of textiles can be enhanced in a multitude of ways, including through the incorporation of core-sheath yarns ^30^, the use multiple layers of dense woven fabrics ^31^, the employment of hard outer layers to blunt the knife ^32,33^, by designing textiles that increase friction between the knife and the yarns, and by increasing inter-fibre friction ^34–37^. It is also fundamentally understood that stab-resistant textiles should have the capacity to absorb energy by causing knife deflection, prior to penetration ^38–40^. High transverse friction exerted on the knife blade can prevent a run-through ^41^, while maintaining a degree of comfort under varied climatic conditions ^42–45^. Similarly to many stab resistant textiles, silk fibre consists of a fibroin core made up of microfilaments and coated by sericin ^46^, which resembles essentially, a core-sheath yarn structure. Additionally, silk cocoons are multilayered ^2,6,8–11^, with the outer layers typically of higher toughness than the inner ones ^12,17^, also ensuring effective thermoregulation for the residing pupae ^1,14,16,47,48^. Based on these notable similarities, we hypothesise that cocoons have evolved stab (penetration and perforation) resistant properties and characteristics.

## MATERIALS AND METHODS

### Material Selection

*Bombyx mori* cocoons were procured from the Kabondo Silk Factory and Marketplace, Kisumu, Kenya. 80 cocoons with uniform shapes and diameters large enough to undergo stab testing were selected. The 80 cocoons were then separated into 2 groups of 40 for testing (i) using a dynamic drop stab test method, and (ii) using an Instron 3369 for static stab testing. Each group was divided into 2 sets of 20 cocoons to separate and test (i) the entire cocoons (EC) and (ii) the cocoon wall segments (CWS). Figure 1 provides an illustration of the experimental design. The entire cocoon (EC) is uncut and simply mounted on the clay for stab testing Figure 2(a), while the cocoon wall segment (CWS) is prepared by cutting the dome-shaped ends of the cocoons parallel to the equator, after which the middle cylindrical section was cut in the meridional direction to obtain a rectangular sheet. The CWS sheets were then mounted between metallic clamps to secure the sample for stab testing, Figure 2(b). Equatorial and meridional directions of the cocoon are denoted with E and M, respectively. Measurements of the EC were taken using Vernier calipers (Hilka Tools Ltd., Chesington, UK), while the thickness of CWS was quantified using a digital screw gauge (Hilka Tools Ltd., Chesington, UK). The apparent density of the cocoon wall was calculated by 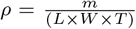, where *ρ* is the density, *m* is the dry weight of the cocoon walls measured using an electronic weighing scale (Ohaus - Cadmus Ltd, Chelmsford, UK) and *L, W*, and *T* represent the length, width and the thickness of the CWS specimen. All specimens were conditioned at a temperature of 21 ^*°*^ and a relative humidity of (55%), Table 1.

**Table 1:** *Bombyx mori* silk cocoon: dimensional measurements and measured densities (includes the range and the mean ± the standard deviation, n = 20).

|  | EC Diameter |  | CWS Measurements |  | Wall Thickness<br>(mm) | Wall Apparent<br>Density<br>(kg/m<sup>3</sup>) |
| --- | --- | --- | --- | --- | --- | --- |
|  | Meridional<br>(mm) | Equatorial<br>(mm) | Meridional<br>(mm) | Equatorial<br>(mm) |  |  |
| Range | 24.3-33.0 | 15.6-19.9 | 13.5-22.0 | 54.6-63.1 | 0.49-0.92 | 283-401 |
| Mean | 29.7 ± 2.0 | 17.4 ± 0.9 | 16.9 ± 2.3 | 57.01 ± 2.6 | 0.68 ± 0.10 | 347 ± 31 |

**Figure 1:**
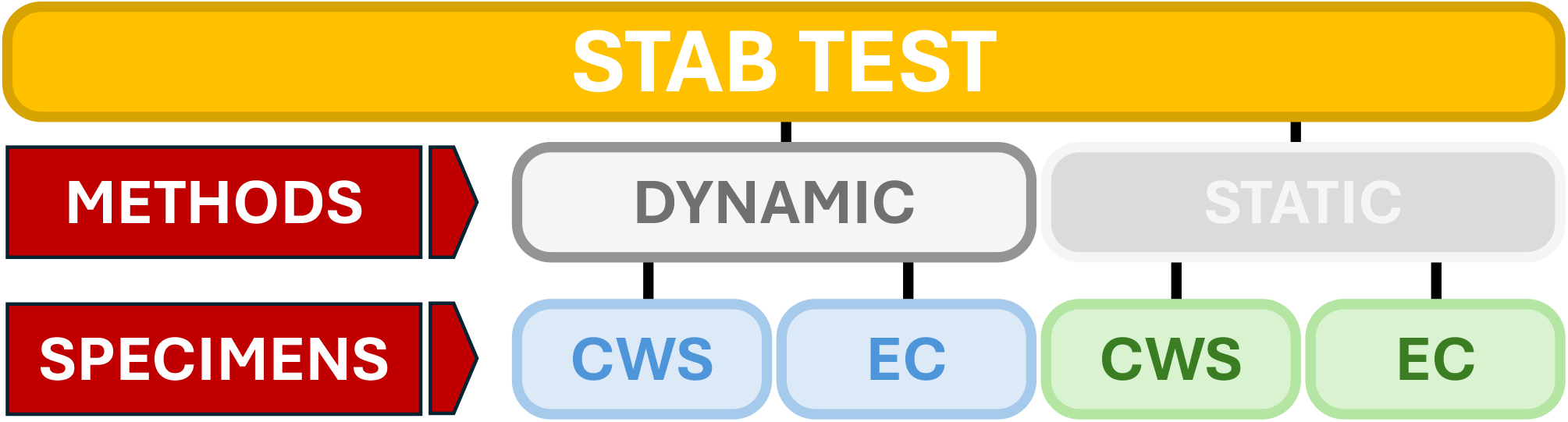
Illustrated experimental design for stab testing showing two methodologies (dynamic and static) with two specimen types tested in each: cocoon wall segment (CWS) and entire cocoon (EC)

**Figure 2:**
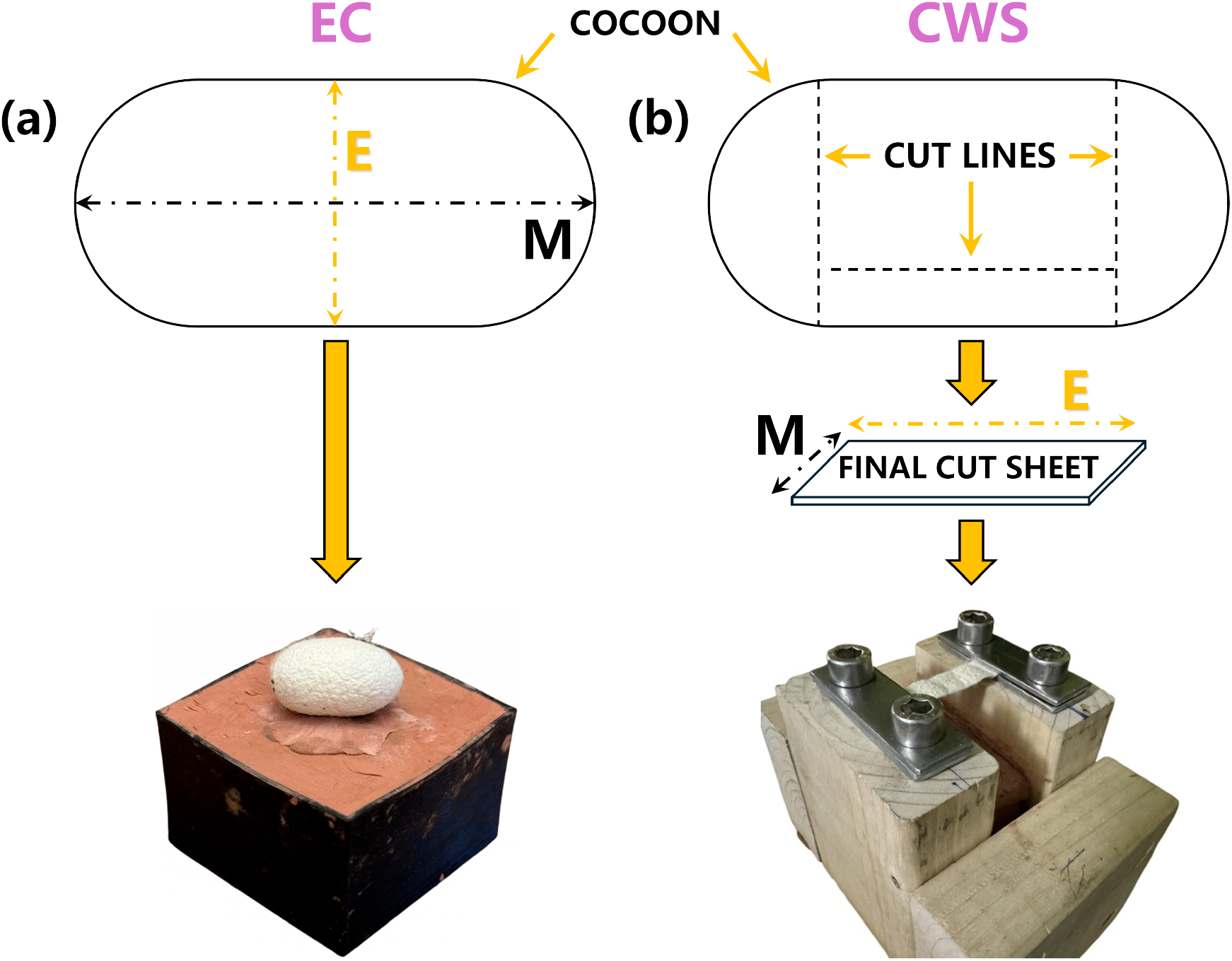
Examples of specimen preparation and mounting: (a) *Bombyx mori* entire cocoon (EC) is uncut and simply mounted on the clay for stab testing and (b) the cocoon wall segment (CWS) is prepared by cutting both poles off a cocoon after which the cut is made along the meridional axis to create a rectangular sheet that is mounted between metallic clamps to secure the sample for stab testing.

### Equipment

A HOSDB P1/B knife blade in all stab tests. The blade has double bevel edges, is 2 mm thick, and 15 mm wide. The knife blade angle is 15^*°*^*±* 0.3^*°*^, with a length of 100 *±* 2 mm, Figure 3(a). A custom fixture was manufactured to hold the CWS flat and prevent slippage during testing, ensuring the tests were reproducible, Figure 3(b). Simultaneously, a 45 *×* 45 *×* 35 mm^3^ box was 3D printed to facilitate the testing of EC specimens. Both fixtures were filled with standard NSP Chavant Clay (Crea Fx, Firenze, Italy), which prevents the knife point from dulling during testing, Figure 3(c). These fixtures along with the HOSDB knife were used for both static testing in an Instron 3369 (Instron Ltd, High Wycombe, UK) and for dynamic testing using the drop test method. Inspired by the National Institute of Justice (NIJ) standard 0115^49^ and the Home Office Body Armour standard 2017^50^, the drop test equipment was designed and manufactured in the lab, Figure 3(d). The equipment comprised a steel frame on which additional components were mounted. The stab sabot assembly consisted of a hollow PVC tube with a knife mount (containing a knife) at one end and a bracket at the other. The bracket was equipped with bearings to facilitate motion over the guide rail. The guide rail was attached to an aluminum extrusion, which was itself mounted onto the steel frame. A drop pin was used to secure the sabot assembly (sabot, knife mount, knife, and sliding bracket) above the testing specimens at the required height. The entire assembly could then be released and dropped onto the specimen by removing the drop pin.

**Figure 3:**
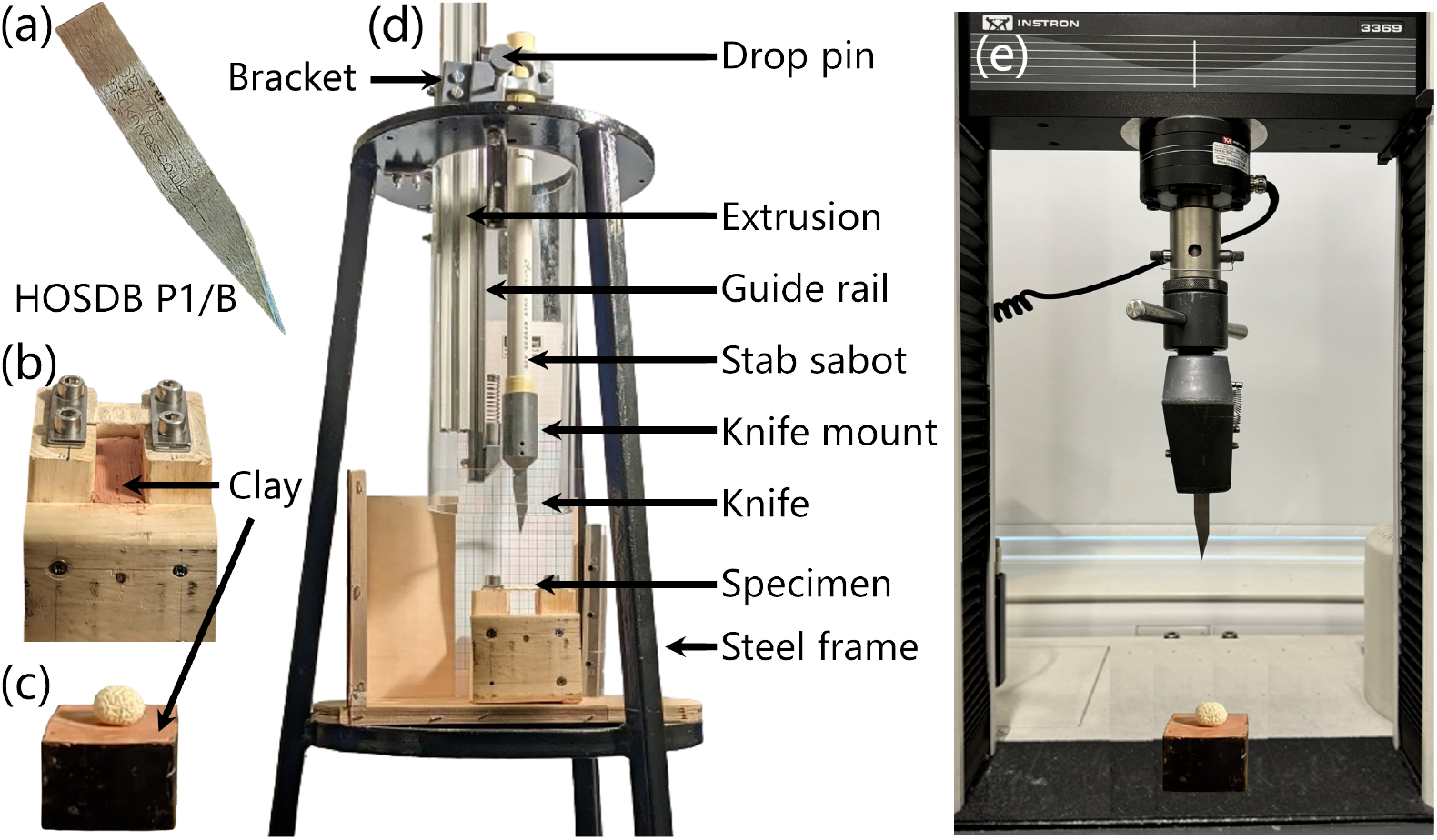
Knife stab experimental setup (a) HOSDB P1/B knife (b) fixture design to hold CWS (c) Container design for testing EC (d) Drop test set up for dynamic testing and (e) Instron 3369 set up for static testing.

### Dynamic stab tests: drop test method

Knife height was adjusted for CWS specimens to ensure that the knife would stab through the specimen but would stop short of impaction with the clay. This ensured that the stab energy was absorbed by only the test specimen, and not by the underlying clay. The fixture, with CWS clamped at both ends, was positioned such that the knife would strike the centre of the specimen, parallel with the equatorial axis, and the knife was released by removing the drop pin. Time and distance was estimated to quantify the energy changes during the process of stabbing. A Chronos 1.4 (Krontech Technologies, British Columbia, Canada) high-speed monochrome camera was used to film the experiment at 1057 frames per second. A one-centimeter square grid was placed behind the stabbing assembly, as shown in Figure 3d, to scale the output and measure the distance travelled in each frame analysed. The entire run of the sabot assembly was divided into four distinct phases which are determined by the position, *P*, of the knife tip: (1) Acceleration (from *P*_0_ to *P*_1_), (2) Deflection (from *P*_1_ to *P*_2_), (3) Penetration (from *P*_2_ to *P*_3_), and (4) Perforation (from *P*_3_ to *P*_4_), Figure 4(a). Here, *P*_0_, *P*_1_, *P*_2_, *P*_3_, and *P*_4_ represent the position of knife tip as follows:

**Figure 4:**
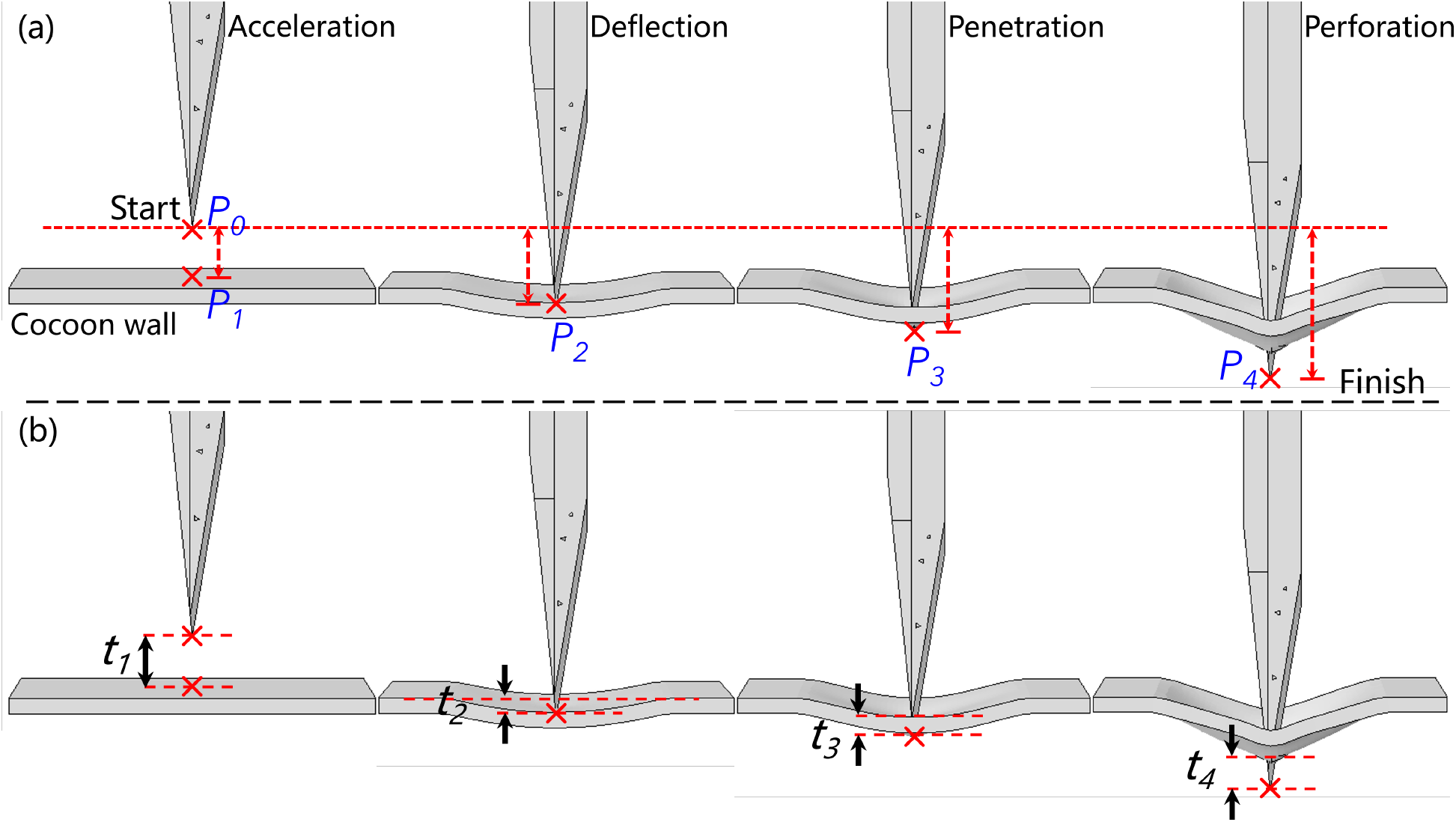
Steps during the process of dynamic stab testing (a) separated into the four stages of acceleration, deflection, penetration and perforation, and (b) time intervals over these four stages.

- *P*_0_: The knife tip is stationary above the material
- *P*_1_: The knife tip makes initial contact with the uppermost surface of the material
- *P*_2_: The knife tip is in contact with the uppermost surface of the material, which reaches its limit of deflection (without the knife penetrating)
- *P*_3_: The initial point at which the knife tip protrudes from the lowermost face of the material (post-penetration)
- *P*_4_: The position of the knife tip after the knife stops moving (post-perforation of the material)

Time intervals, *t*_*n*_ correspond to the different stages of acceleration (*t*_1_), deflection (*t*_2_), penetration (*t*_3_), and perforation (*t*_4_). These time intervals align with the knife tip travel, from *P*_0_ to *P*_1_, *P*_1_ to *P*_2_, *P*_2_ to *P*_3_, and *P*_3_ to *P*_4_, respectively, Figure 4(b). Knife displacements, *h*_*n*_, over each stage are thence represented as *h*_1_, *h*_2_, *h*_3_ and *h*_4_, representing acceleration, deflection, penetration and perforation stages respectively. The velocities, over each stage, are represented by *v*_1_, *v*_2_, *v*_3_, and *v*_4_ over each of the stages, representing acceleration, deflection, penetration and perforation stages respectively. These are calculated in accordance with Equation (1).

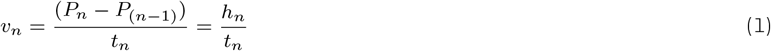

Finally, the corresponding changes in kinetic energy of the sabot assembly, Δ*KE*_*n*_, were calculated using Equation (2) over the different stages of acceleration (Δ*KE*_1_), deflection (Δ*KE*_2_), penetration (Δ*KE*_3_), and perforation (Δ*KE*_4_). These align with the knife tip travel, from *P*_0_ to *P*_1_, *P*_1_ to *P*_2_, *P*_2_ to *P*_3_, and *P*_3_ to *P*_4_, respectively, and *m*_*s*_ defines the mass of the sabot assembly.

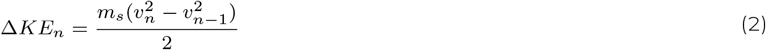

The drop tests for EC specimens followed the same procedure as with the CWS specimens, except with the full uncut cocoons placed simply over the clay-filled container. Since the knife tip was no longer visible after penetrating the uppermost wall of the cocoon, the energy calculations made for the CWS specimens could not be made for the EC specimens. However, stabbing the entire cocoon provided us with the opportunity to analyse dimensional changes through image analysis.

### Static stab tests: Instron 3369

The stab resistance of both CWS and EC specimens were evaluated using an Instron 3369 (Instron, High Wycombe, UK), cf. Figure 3(e). The knife was secured in the top grip of the Instron, equipped with a 1 kN load cell. To ensure uniformity during wall stabbing, the knife width was arranged parallel to the equatorial direction of the CWS specimen and its tip was aligned to the centre of mass of the specimen. For testing EC specimens, the cocoon was placed on the clay-filled container, with the knife positioned at the intersection of the meridional and equatorial axes, with the blade aligned parallel to the meridional axis, again to maximise the available material for the knife to stab through. Static testing was conducted at a rate of 20 mm/min. The test was recorded using similarly to the drop test, and complete experimental setup showing the position of the camera is illustrated in Figure 5. In all cases, the camera was positioned in line with the sample, so that lateral movement of both EC and CWS samples could be captured accurately.

**Figure 5:**
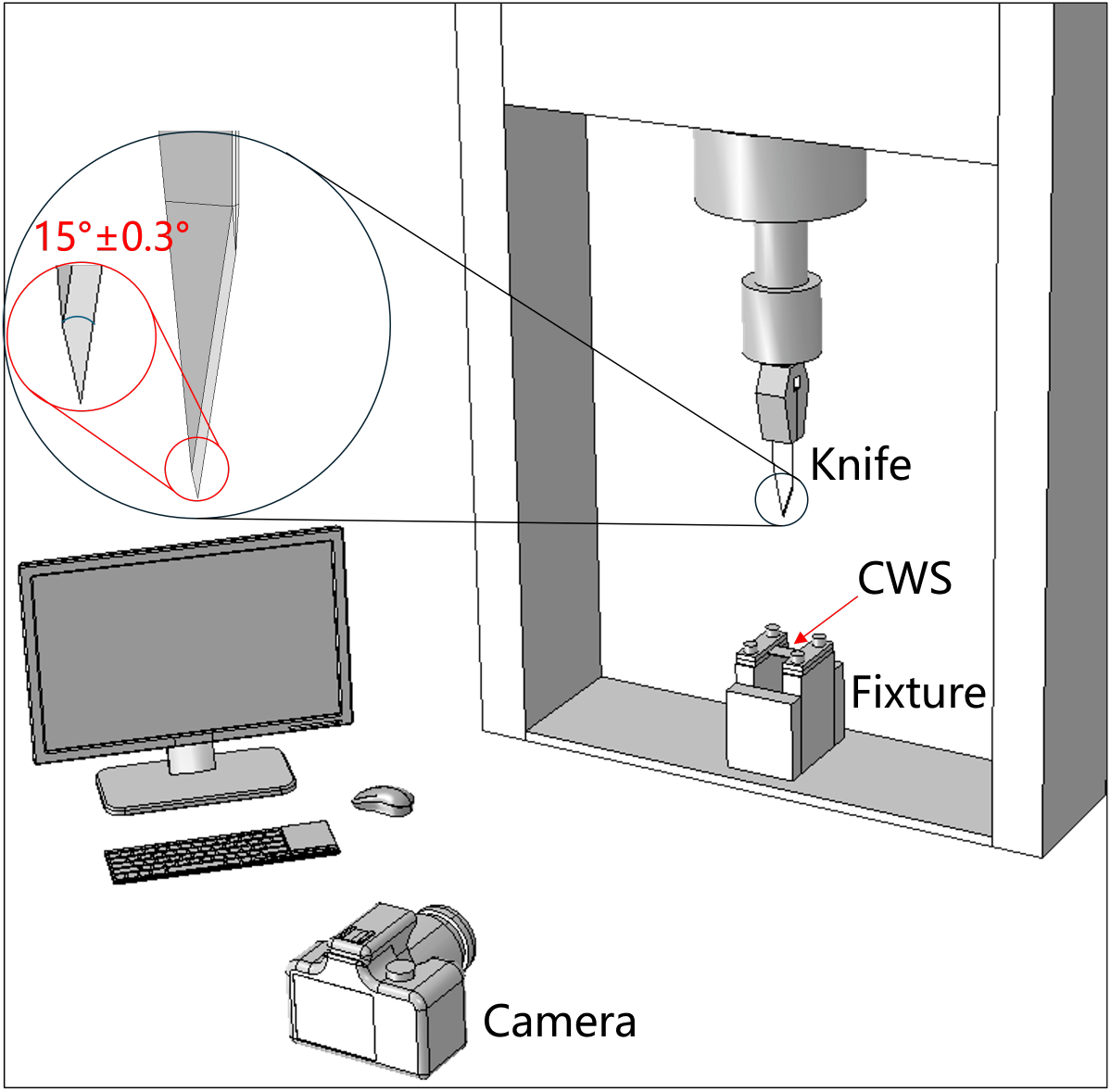
Experimental arrangement for the stab testing of CWS and recording camera

### Static tensile testing

Cocoon wall specimens (CWS) were tested in tension to determine the tensile properties of the material. All CWS were prepared with a uniform width of 12 mm. The gauge length was set at 10 mm in each sample and each specimen was tested in the equatorial axis of the cocoon at a crosshead speed of 10 mm/min. The rationale for testing exclusively in one direction was that the properties and characteristics of the *Bombyx mori* cocoon have been documented to exhibit similarity across various directions ^4,5^.

### Finite element modelling

We use an explicit dynamic finite element method to simulate and analyse cocoon deformation from a knife stab, using the commercial software ABAQUS/CAE 2021 (Dassault Systèmes Simulia Corp., Johnston, RI, USA). The geometry of the cocoon was designed as a revolving shell, exhibiting symmetry around its meridional axis. S4R shell elements were used to discretise the cocoon geometry, with the dimensions specified in Table 1. S4R are 4-node, quadrilateral, stress-displacement shell elements. These have the option for reduced integration and importantly, a large-strain formulation appropriate for the types of expected cocoon deformations. The final mesh comprised 16,424 S4R elements. The knife geometry was defined as an undeformable solid (rigid body), and the dimensions were based on the HOSDB P1/B standard knife specifications ^50^. An elastic-perfectly plastic model was employed to simulate cocoon damage, which describes the strain response to applied stress, Equations (3) and (4) ^51^.

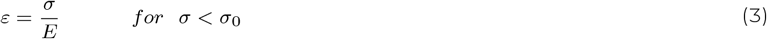

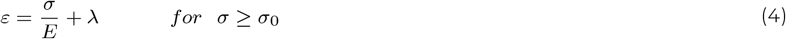

Here, *ε* is the strain, *σ* is the stress, *E* is the Young^*′*^s modulus, *σ*_0_ is the yield stress, and *λ* is a scalar (Lame’s coefficient) greater than zero. Physical and mechanical properties input into the model were obtained through measurements of the dimensions and densities, as well as through tensile testing, which we provide in Table 1 and Table 2, respectively. The bottom of the cocoon had a fixed boundary condition, constraining all degrees of freedom (Encastre boundary). All degrees of freedom of the knife were constrained except in terms of its translation at 5 m/s in the direction of the cocoon.

**Table 2:**
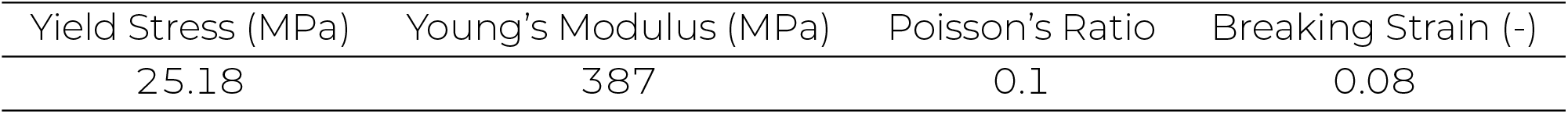
Elastic-perfectly plastic model parameters derived from our tensile testing, used as input parameters in the finite element model.

### Image analysis

Dimensional changes of the EC specimens during the stab test were analysed using the image analysis software Fiji ^52^. Changes in the meridional and equatorial dimensions were estimated using the line measurement tool. Accurately measuring changes in the cocoon volume during the stabbing process was not possible since only one camera was available. As such, changes in the face projection area of the cocoon were studied as a proxy for volume changes. Images captured at various stages of the stabbing process were thresholded to estimate area changes accurately, allowing for a detailed understanding of how the cocoon dimensions were affected throughout the stabbing process.

### RESULTS AND DISCUSSION

Using a coupled oven drying-weighing method, EC and CWS were found to have moisture contents of 4.9 % and 4.1 %, respectively. Following the degumming procedure outlined in ^10^, the fibre content in the native cocoons was determined to be 74 % by weight, with a standard deviation of 1.73 %. Figure 6 shows the stress-strain curves of five cocoon specimens tested in tension. The tensile strength, strain at max strength, and Young’s modulus of the cocoon material were found to be 25.2 *±* 2.3 MPa, 0.078 *±* 0.009, and 387.2 *±* 27.9 MPa, respectively. These values lie within the ranges of previously values reported for tensile strength from 13 *±* 0.25 to 54 *±* 11 MPa ^5,10,12,13,53^, for tensile strain from 0.05 to 0.2^13,55^, and for tensile modulus 300 *±* 20 to 586 *±* 111 MPa ^5,10,12,13,53^, respectively. We measured the toughness of *B. mori* cocoons at 1.14 *±* 0.27 MJ *mm*^−3^, which is in line with earlier published toughness values, reported at 1.1 *±* 0.01 MJ *mm*^−3^ ^13^. The mechanical properties measured in our work have an expected range in terms of variance ^54^. The physical properties measured in both the EC and CWS specimens have been summarised in Table 1.

**Figure 6:**
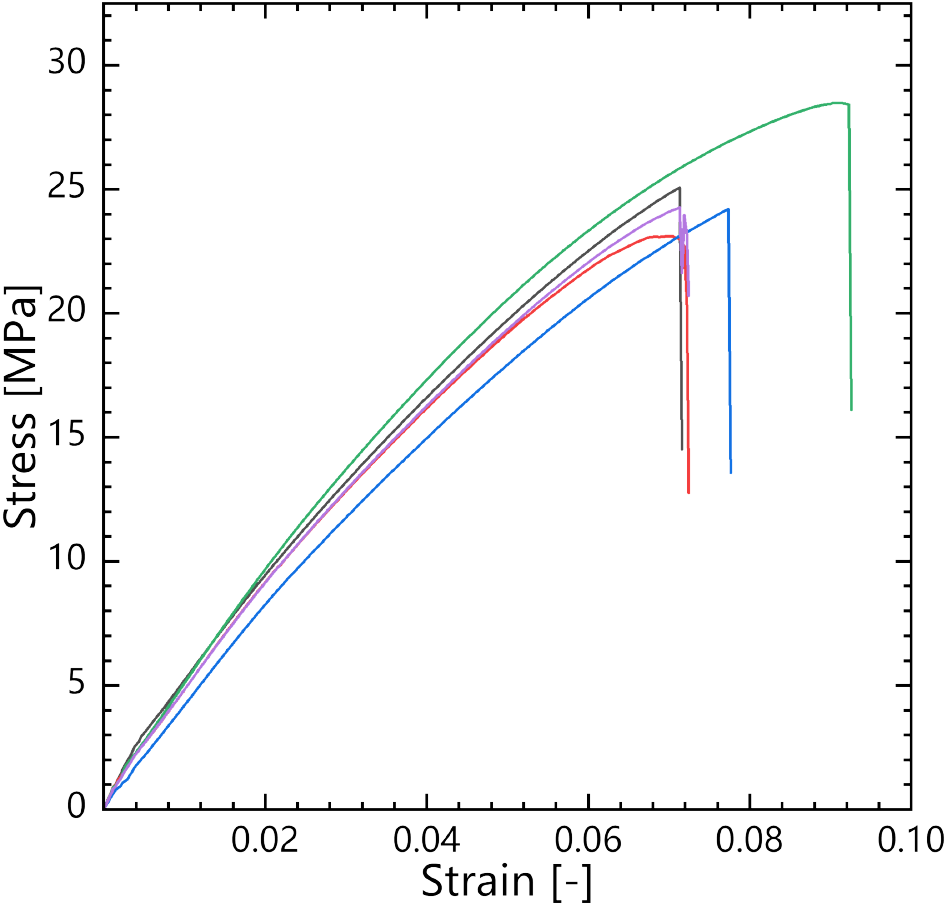
Stress-strain curves of *Bombyx mori* cocoon wall specimen (n = 5).

### Dynamic stab testing: drop test method

Dimensional changes during dynamic stabbing were analysed for all EC specimens (n=20). Captured videos were split into frames which were then processed using Fiji software. The mean cocoon size decreased from 33.7 *±* 2.1 mm to 33.2 *±* 2.1 mm meridionally, and from 22.3 *±* 1.4 mm to 19.3 *±* 1.4 mm equatorially, Figure 7(a-b). Corresponding strains in the meridional and equatorial dimensions were of the order of 0.01 *±* 0.008 and 0.13 *±* 0.04, respectively. Additionally, changes in the apparent area were examined using 2D projections of the cocoon face, Figure 7(e). It was noted that the mean apparent area decreased from 607 *±* 48 mm^2^ to 568 *±* 35 mm^2^, Figure 7(d). To check the statistical significance, one way ANOVA was performed. A p-value of 0.46 (at *α* = 0.05) implied that the change in the meridional direction was not significant. Changes in the equatorial dimension, differences between the meridional to equatorial strains, and the changes to the apparent area were found to be statistically significant since p < 0.001, p < 0.001 and p < 0.01, respectively, when comparing these before and after stabbing. Box plots in Figure 7(a-d) present the full distributions of data about the mean and median values in each data set. Apparent areal changes in the cocoon were examined for knife penetration. The apparent area at any point of stabbing (*A*) was normalised by dividing it by the underformed apparent cocoon area (*A*_0_) and this is plotted against knife displacement, Figure 7(e). A non-linearly proportional decrease in the normalised apparent area is noticeable as knife penetration increases linearly. The fitted line in this figure starts to become asymptotic with the knife displacement axis at ca. 12 mm knife displacement. In Figure 7(e), the finite element model output is compared against the experimental measurements and observably follows a similar trajectory, closely mapping the experimental dimensional changes as a function of knife displacement, especially where *h*(*t*) ≤ 6 mm. Figure 8 illustrates the simulated dimensional changes, obtained from ABAQUS/CAE model outputs.

**Figure 7:**
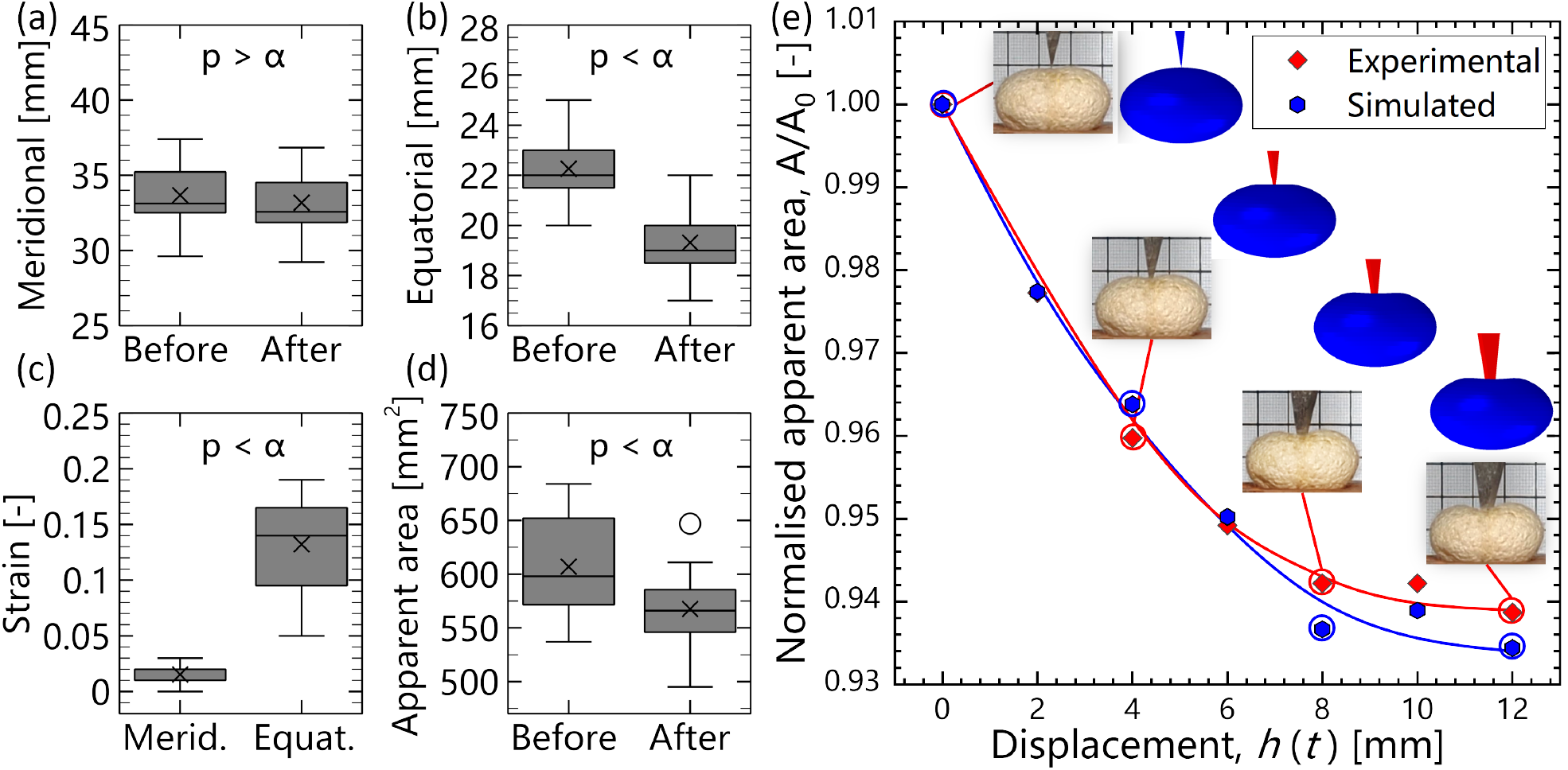
Measured dimensional changes of the cocoon before and after stabbing in (a) the meridional direction and (b) equatorial direction. Changes in strain due to stabbing in meridional and equatorial dimensions are shown in (c). In (d), changes to the cocoon apparent area (i.e. a 2D projection of a 3D body) are shown, and (e) shows changes in the normalised apparent area against knife displacement, comparing both the experimental samples and the finite element model output. Error bars represent the standard error about the arithmetic mean (n = 20).

**Figure 8:**
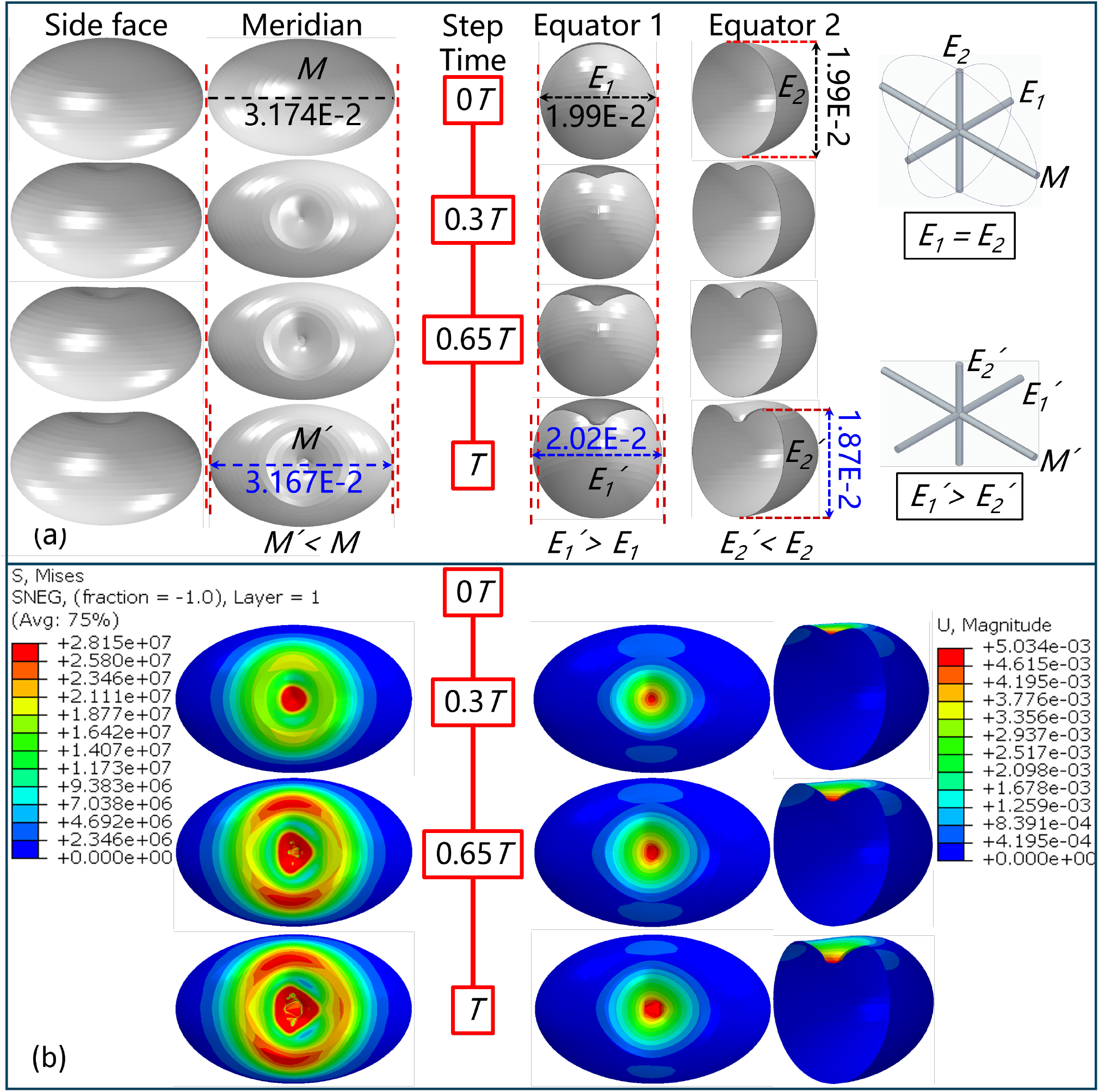
Simulated dimensional changes in the cocoon showing (a) changes in side face, meridional, and two equatorial axis, (b) distribution of stress over the area under the knife point and magnitudes of displacement over the cocoon top section. Here, *T* is the total simulation time.

*M* defines the measure of the meridional axis while *E*_1_ and *E*_2_ define the two equatorial axes, which are both equal and perpen-dicular to one another. When a stabbing force is applied in line with *E*_2_, the cocoon contracts along *M* and expands along *E*_1_. The expansion of the cocoon in *E*_1_ when compressed along *E*_2_, may be attributable to its ellipsoidal geometry, which when compressed in one dimension transfers the applied stress radially, causing expansion ^15^. This can also be seen as a cocoon level Poisson’s effect, as the cocoon expands perpendicularly under an applied compression ^16^. Since the finite element model deformations aligned well with the experimentally measured deformations (cf. Figure 7(e)), and they provide more discrete and extractable numerical values for deformation than those measured from the video frames, we used the model outputs to determine the Poisson’s ratios of the cocoon. We find that the cocoon displays a unique anisotropic behaviour in response to knife stabbing, with Poissons ratios of the order of 0.25 and −0.05 along *E*_1_ and *M*, respectively. An auxetic response was observed, with a negative Poissons ratio (−0.05) is noted to occur along the meridional axis *M*. These variations in Poisson’s ratio along two orthogonal axes indicate that the cocoon may be geometrically optimised to maximise protection through controlled deformation as the expansion in *E*_1_ coupled to a contraction in *M* helps conserve space in *E*_2_.

Both the von Mises stresses, *S*, and cocoon deformations, *U*, are rendered in Figure 8(b), which shows sequential images taken at approximately equal intervals throughout the simulation. The highest stress concentrations appear localised at this tip and initially (0.3*T*) dissipate radially from the contact point. As the stresses evolve from the knife tip (> 0.65*T*), the stresses adopt a triangular profile, following the contours of the knife cross sectional geometry. This geometry leads to an uneven stress distribution, with elevated stresses around the base angles of the triangular blade profile. The loading applied by the knife creates a deformation concavity in the cocoon material surrounding the contact point, with the highest point of deformation arising directly under the knife tip. During the formation of the concavity, the uppermost regions of the cocoon both flex and tension during deformation, resulting in the development of high stress concentrations. In addition to this, as reported by Chen et al. ^26^, the combination of an ellipsoidal geometry, a fibrous network, a laminated structure, and the graded material properties of the cocoon are likely contributors to mechanical resistance.

Energy changes in the sabot assembly were analysed for tests conducted on the CWS from the starting position *P*_0_ to the endpoint *P*_4_, Figure 9. The average displacement of the knife tip from *P*_0_ to *P*_4_ was 0.0865 *±* 0.0062 m, and the time taken was 0.116 *±* 0.012 seconds. The sabot assembly achieved a peak kinetic energy of 0.291 J of which 0.146 *±* 0.019 J was during the acceleration phase while 0.145 *±* 0.058 J was gained over the deflection phase. Collectively, this region could be regarded as a region of energy acquisition. Conversely, the sabot assembly dissipated 0.277 *±* 0.055 J and 0.014 *±* 0.008 J over a region of energy dissipation, which consisted of both penetration and perforation phases, respectively, Figure 9. Notably, 95% of the absorbed energy was dissipated during penetration. The remaining 5% of the energy facilitated knife perforation through the material, ultimately leading to cessation in movement. Table 3 provides a detailed account of the displacements among predefined positions, corresponding times and velocities, and the associated kinetic energy changes.

**Table 3:** Energy changes in sabot assembly during the stabbing.

|  |  | Displacement<br>(dm) | Time<br>(ms) | Final Velocity<br>(dm.s<sup>-1</sup>) | Kinetic Energy Change<br>(mJ) |
| --- | --- | --- | --- | --- | --- |
| Start point | $P_0$ | 0 | 0 | 0 | 0 |
| Acceleration | $P_0-P_1$ | $0.640 \pm 0.02$ | $121 \pm 4.8$ | $8.3 \pm 0.5$ | $146 \pm 19$ |
| Deflection | $P_1-P_2$ | $0.064 \pm 0.02$ | $5.5 \pm 1.6$ | $11.6 \pm 1.1$ | $145 \pm 58$ |
| Penetration | $P_2-P_3$ | $0.006 \pm 0.001$ | $2.7 \pm 0.3$ | $2.5 \pm 0.6$ | $-277 \pm 55$ |
| Perforation | $P_3-P_4$ | $0.155 \pm 0.027$ | $37.2 \pm 5.5$ | 0 | $-14 \pm 8$ |

**Figure 9:**
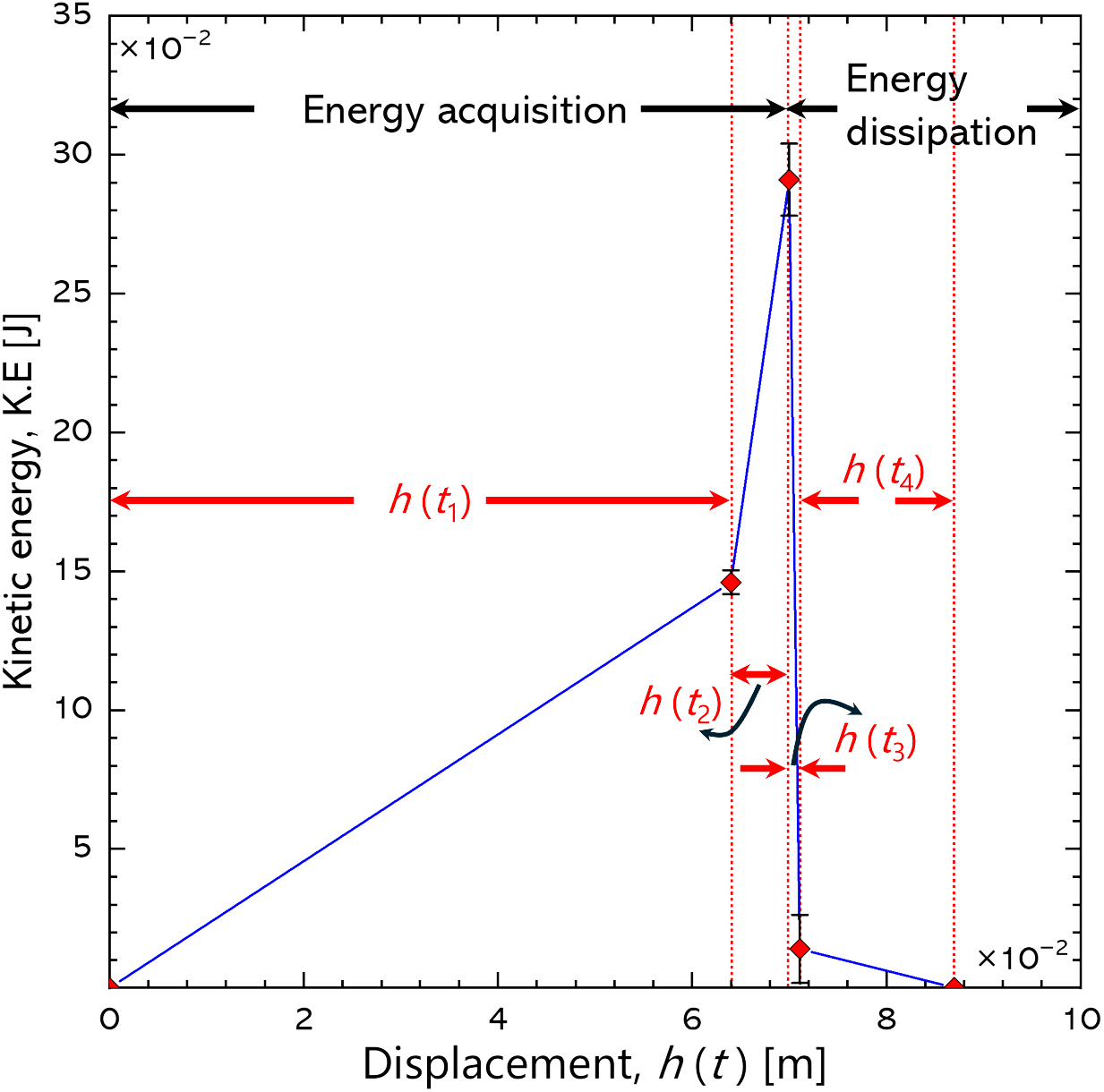
Energy changes in stabbing sabot assembly. Error bars represent the standard error about the means.

### Static stab tests: Instron 3369

Mean stabbing forces for the EC and the CWS were of the order of 5.65*±*1.07 N and 5.42*±*1.20 N, respectively, Figure 10. The median stab forces for the same were of the order of 5.64 N and 5.63 N, respectively. The stab force data ranged from 4.03 to 7.74 N for EC and 3.5 to 8.18 N for CWS specimens. One-way ANOVA was used statsitically determine the level of difference between the stab force datasets of EC and CWS. A p-value of 0.86 was obtained and is considerably higher than the significance threshold (*α* =0.05), while the computed F-statistic (F = 0.03) was lower than the critical F-value of 4.1, table 4. As such, we conclude that there is no discernible difference between the two datasets.

**Table 4:** One-way ANOVA comparing stab force between the EC and the CWS sample sets. Here, ss is the sum of squares, d is the degree of freedom, MS is the mean sum of squares, F is the F-statistics and F crit is the F-critical value.

| Source of Variation | SS | df | MS | F | P-value | F crit |
| --- | --- | --- | --- | --- | --- | --- |
| Between Groups | 0.04 | 1 | 0.04 | 0.031 | 0.86 | 4.1 |
| Within Groups | 47.85 | 38 | 1.26 |  |  |  |
| Total | 47.89 | 39 |  |  |  |  |

**Figure 10:**
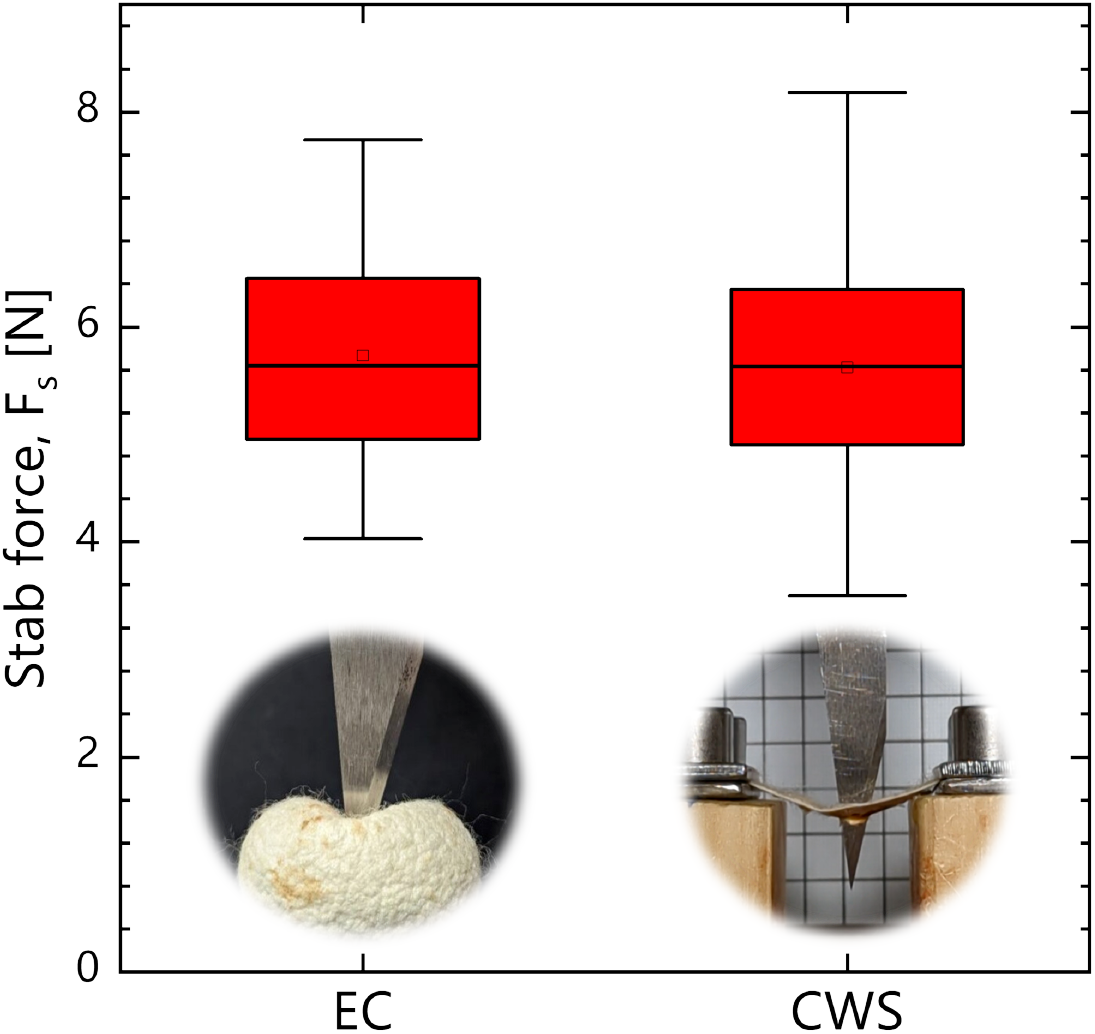
Comparison of stab force: entire cocoon (EC) vs cocoon wall specimen (CWS). Error bars represent the standard deviation of the means. One way ANOVA: p = 0.86 for *α* = 0.05 (no difference between measured stab forces of EC and CWS).

The static stab force is plotted in Figure 11 against knife displacement for a representative (a) EC specimen and (b) CWS specimen. These graphs highlight key differences in material behaviour, which are in turn related to specimen geometry. Observing the curve for the EC specimen in Figure 11(a), we note the force-displacement curve exhibits initial linear elastic behaviour. This is in contrast to the force-displacement curve of the CWS sample, where the stab force increases nonlinearly at a higher rate of change as a function of a linear increase in knife displacement. When this curve reaches a displacement close to 2mm, the curve then exhibits a linear elastic behaviour. The reason for this is that CWS specimens are cut from a curved cocoon, and as such, can never become truly flat test coupons. Rather, there will always remain curvature in the CWS, which leaves a degree of slack in the specimen that has to be overcome through knife movement before the material becomes sufficiently taut and is able to deform elastically. Although both EC and CWS specimens ruptured at comparable stabbing forces, the first fracture peak occurs at a knife displacement of 1.3 mm and 3.2 mm for the EC and CWS specimens, respectively. This indicates that the stab displacement for the EC was two-fifths of the CWS. Upon reaching the final rupture point, both EC and CWS specimens exhibit a catastrophic drop in the stabbing force, which is related to initial penetration. The stab force continues to increase after this drop in force as the knife continues to penetrating and perforate through the cocoon material. This is related to the knife geometry, which increases in size from tip, increasing resistance the area of material needed to penetration and perforation, as a function of knife movement. The stab force reaches its maximum at above 8 N in both EC and CWS specimens, Figure 11(a-b).

**Figure 11:**
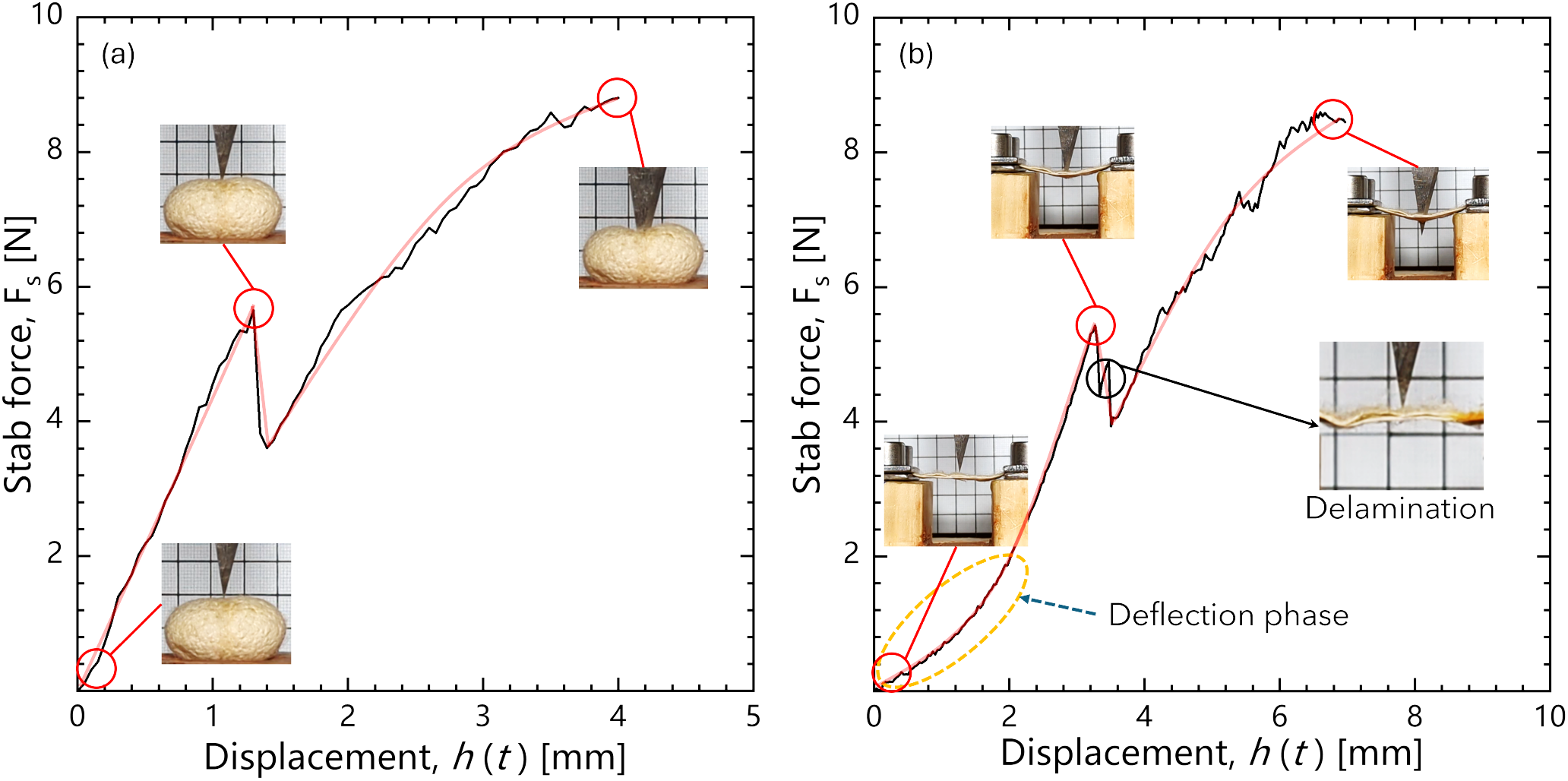
Comparison of stab force against knife displacement (a) EC (b) CWS.

Naturally, a cocoon possesses a curved, multilayered graded wall, with inner layers having a smaller circumference than outer layers. Cutting and attempting to flatten the CWS specimen can induce microfibre buckling, sericin cracking and delamination, as illustrated in Figure 12. This perspective is supported by the literature, where it is stated that failure in cocoon wall material is instigated from sericin cracking at interfaces between layers (interlayers), which is weaker than the intralayer fibre-sericin bonding ^5,55^. Such delaminations may result in a stepped failure behaviour, as observed in the CWS specimens shown in the delamination insert in Figure 11(b). The geometry of the cocoon being a continuum, uncut structure, will also likely mitigate delamination fractures more effectively than a cut sample. Ellipsoids can deform elastically upon impact, storing energy within the structure and releasing it at a critical level proportional to the impact force ^26^.

**Figure 12:**
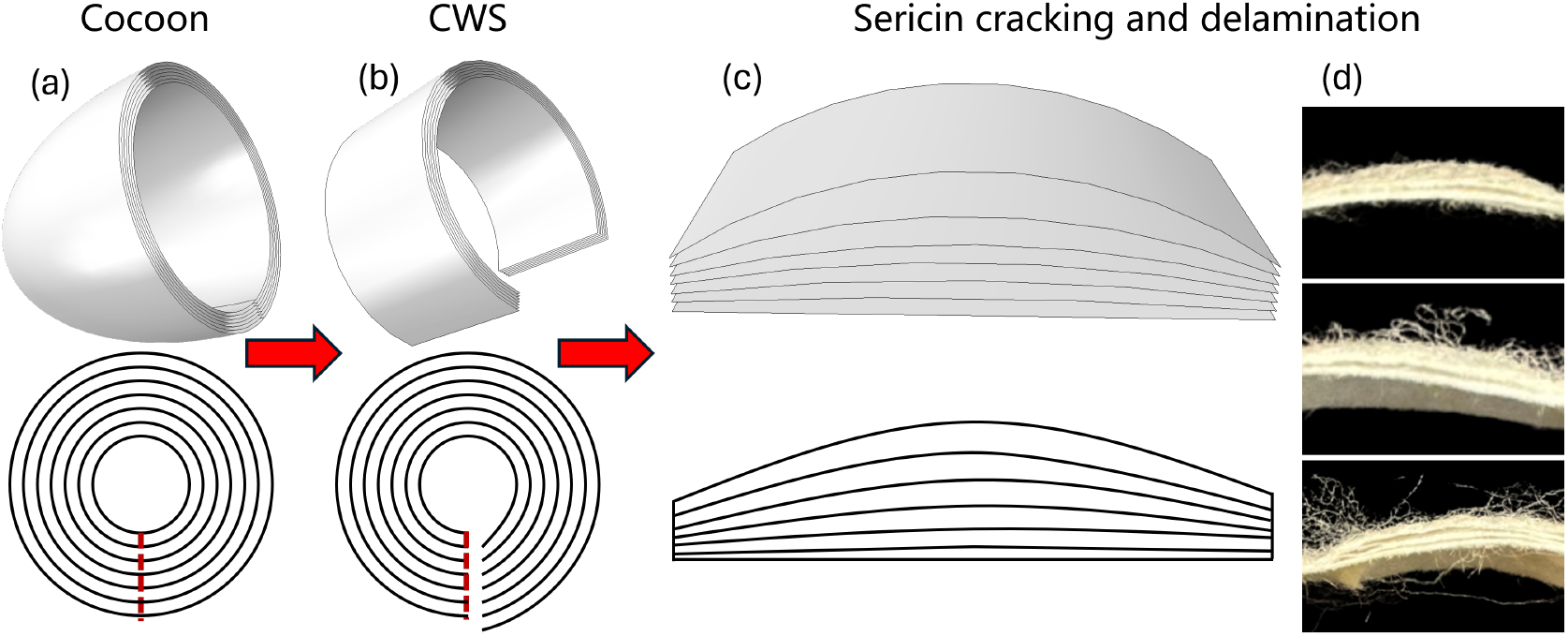
Conversion of a cocoon into a CWS resulting in sericin cracking and delamination: (a) illustration of dome end cuts (b) pre-flattened CWS specimen (c) forced flattening of CWS resulting in sericin cracking and delamination, and (d) evidence of delamination in flattened CWS specimens.

### Stab footprints

On contact, force is transferred from the knife tip to the cocoon material, which absorbs mechanical energy ^56^ by deflection, Figure 13(a-b). When the material reaches a deflection limit, concentrated stresses from under the knife tip dissipate laterally, potentialising interlayer shear and thus delamination failure of the cocoon material ^57^. On penetration, the knife tip disrupts the fibre network, causing fibre stretching, bending and migration. When fibres break, the material ruptures ^9^ and energy dissipates during the penetration phase over the contact area as well as from through thickness penetration, which increases frictional stresses at the knife-material interface, Figure 13(c). After the knife has penetrated the material, the perforation stage begins and the bevelled edges cut through the fibres. The stab force requirement over this stage increases since the knife bevel widens as the stab deepens, Figure 13(d), which in turn continues to increase the contact friction between the knife and cocoon material. This is evidenced by the increased stab force beyond the rupture point (cf. Figure 11), eventually leading to the termination of knife movement since there is no further kinetic energy available to dissipate. Cocoon damage is essentially a coupled failure involving sericin bond breaking and delamination ^26^. Wang et. al ^9^ further note that cocoon wall rupture by needle penetration results from either the frictional force between the needle and the fibre, or from fibre stretching, leading to tensile failure of the fibres, which correlates well with our own observations. Figure 13(e-g) shows the knife stab footprints on the top and bottom faces of the cocoon wall. The knife blade cross-section has a pointed edge at the cutting end and a 2 *mm* width at the opposing end, closely resembling an isosceles triangle. As such, the area of the footprint approximates half the product of the base and the perpendicular height 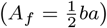. Where *A*_*f*_ denotes the area of the knife foot-print, and *b* and *a* are the footprint base and height, respectively. The stab footprint areas of the tested specimens are summarised in Figure 13(h). The box plot indicates that the mean footprints for the entire cocoons (EC) were of the order of 5.92 *mm*^2^ with a standard deviation of 0.80, while for the cut wall specimens (CWS), the mean footprint was around 8.89 *mm*^2^ with a standard deviation of 1.89. A one-way ANOVA test (at *α* = 0.05) comparing the two datasets yielded p < 0.001, indicating a statistically significant difference between the two sets. The lower mean footprint for the EC suggests that the stabbing knife encountered higher resistance, likely due to the mechanical energy required to deform the large elliptical geometry of the EC, as compared to the that required to deflect a CWS specimen. The lower spread of data in the EC specimens compared to the CWS specimens indicates that footprint damage is more consistent in the EC specimens.

**Figure 13:**
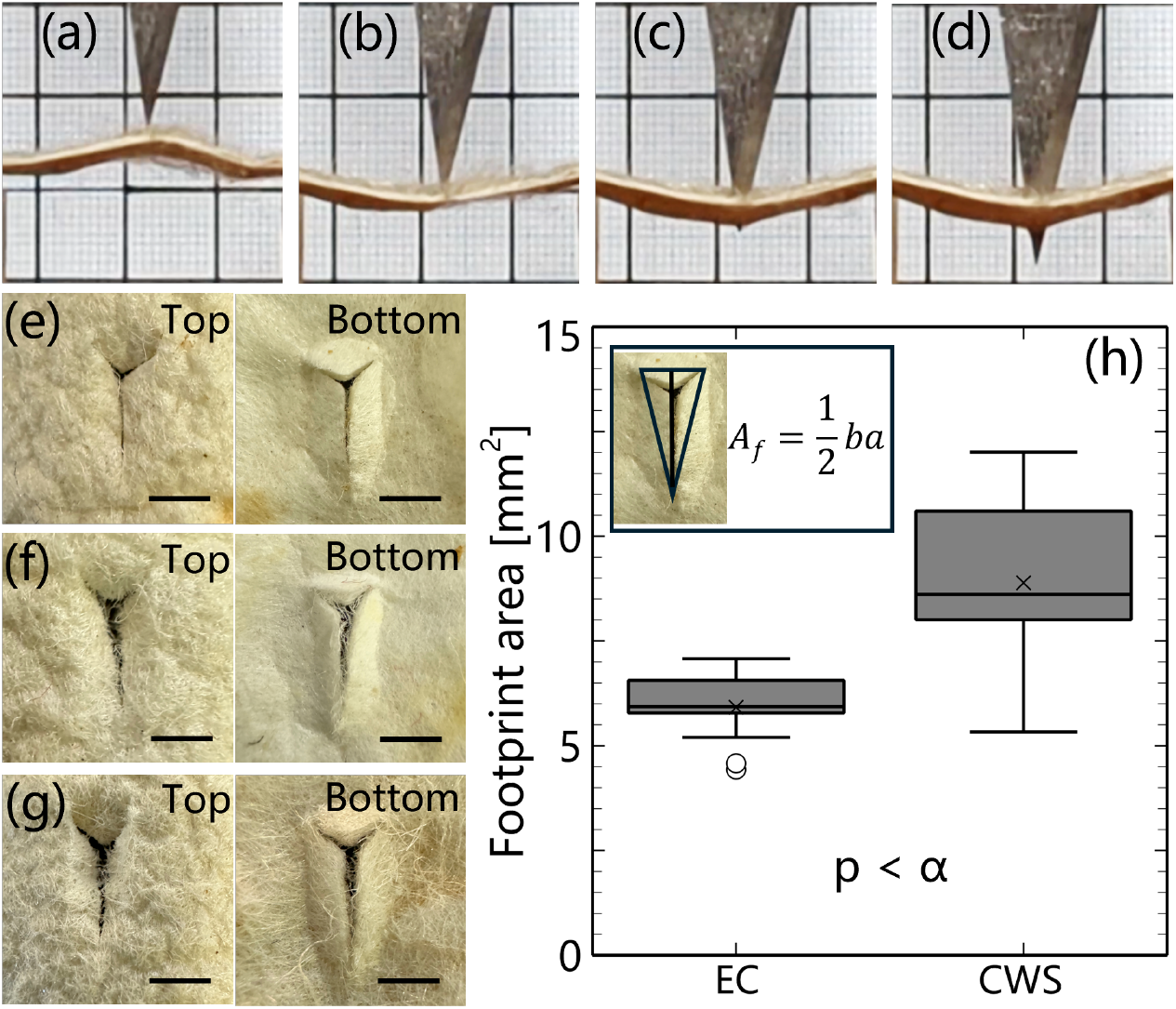
Damage mechanism of the cocoon wall, (a and b) material deflection, (c) penetration, (d) perforation, (e to g) top and bottom knife footprints (scale bar: 2 mm), (h) box plot showing data distribution of stab footprint area of the EC and CWS.One way ANOVA: p *<* 0.001 for *α* = 0.05, indicating there is a significant difference between the stab footprint area of EC and CWS.

## CONCLUSIONS

*Bombyx mori* cocoons were tested for knife stab resistance using both dynamic and static stab testing methods. We tested both entire cocoons (EC), as well as cocoon wall segments (CWS) cut into rectangular quasi-flat sheets. We identify three stages in the stab process, arising from the point of knife contact with either an entire cocoon or a cocoon wall segment The first stage is material deflection, the second stage is knife penetration and the third stage is knife perforation. 95 % of the kinetic energy is lost through the penetration stage. EC specimens exhibited statistically significant differences (p-value < *α*) before and after stabbing in terms of: strain experienced between the meridional and equatorial directions, deformation in the equatorial direction and changes to the apparent area. No significant differences were observed in deformation in the meridional dimension before and after stabbing. Simulated results aligned with experimental graphs when mapping the normalised apparent cocoon area, which in both cases begin to become asymptotic with displacement. Although the mean stab forces needed to penetrate EC (5.65 N) and CWS (5.42 N) were similar, differences were noticed when comparing the force-extension curves. Curves for the EC specimens exhibited less deflection as well as a more uniform increase in stabbing load as compared to the CWS specimens. EC was also noted to exhibit anisotropic behaviour during stabbing, with a positive Poisson’s ratio (0.25) for *E*_1_:*E*_2_ and negative Poisson’s ratio (−0.05), an auxetic response, for *M* :*E*_2_. Knife footprints of EC were notably smaller than the CWS after stabbing, from which we infer that the EC specimens are structurally more stab resistant and that the EC geometry enables improved stab damage tolerance.

## DATA AVAILABILITY

Data for this publication will be made available through Edinburgh DataShare (https://datashare.ed.ac.uk/) and can also be made available from the corresponding author on request.

### ACKNOWLEDGMENTS

The authors wish to thank Mr. Ali Aitzaz from The University of Edinburgh for his gracious help with the high speed filming needed for our research.

## AUTHOR CONTRIBUTIONS

Conceptualization (PA); Data curation (AUR); Formal analysis (AUR, VK, PA); Funding acquisition (AUR); Investigation (AUR); Methodology (AUR, VK, PA); Project administration (PA); Resources (VK, PA); Software (na); Supervision (VK, PA); Validation (AUR, VK, PA); Visu-alisation (AUR); Roles/Writing - original draft (AUR); Writing - review and editing (VK, PA).

## AUTHOR COMPETING INTERESTS

The authors declare no competing interests.

## OPEN ACCESS STATEMENT

For the purpose of open access, the authors have applied a Creative Commons Attribution (CC BY) license to any author accepted manuscript version arising from this submission.

## Notes

### Competing Interest Statement

The authors have declared no competing interest.

